# Subtype specific responses in hKv7.4 and hKv7.5 channels to polyunsaturated fatty acids

**DOI:** 10.1101/2021.08.20.457075

**Authors:** Damon J A Frampton, Johan Nikesjö, Sara I Liin

**Affiliations:** Department of Biomedical and Clinical Sciences, Linköping University, Linköping, Sweden

**Keywords:** docosahexaenoic acid, electrophysiology, electrostatic, KCNQ, lipid, omega 3, two-electrode voltage clamp

## Abstract

The K_V_7.4 and K_V_7.5 subtypes of voltage-gated potassium channels are expressed in several tissues where they play a role in physiological processes such as sound amplification in the cochlea and adjusting vascular smooth muscle tone. Therefore, the mechanisms that regulate K_V_7.4 and K_V_7.5 channel function are of interest. Here, we study the effect of polyunsaturated fatty acids (PUFAs) on human K_V_7.4 and K_V_7.5 channels expressed in *Xenopus* oocytes. We report that K_V_7.5 is activated by PUFAs, which shift the V_50_ of the conductance *versus* voltage (G(V)) curve towards more negative voltages. This response depends on the charge of the head group as an uncharged PUFA analogue has no effect and a positively charged PUFA analogue induces positive V_50_ shifts. In contrast, we find that the K_V_7.4 channel is inhibited by PUFAs, which shift V_50_ towards more positive voltages. No effect on V_50_ of K_V_7.4 is observed by an uncharged or a positively charged PUFA analogue. Oocytes co-expressing K_V_7.4 and K_V_7.5 display an intermediate response to PUFAs. Altogether, the K_V_7.5 channel’s response to PUFAs is like that previously observed in K_V_7.1-7.3 channels, whereas the K_V_7.4 channel response is opposite, revealing subtype specific responses to PUFAs.

## Introduction

The K_V_7 family of voltage-gated potassium channels are expressed in several tissues where they serve to attenuate excitability by conducting an outward K^+^ current. The five members of the family, K_V_7.1-K_V_7.5 (encoded by KCNQ1-KCNQ5 genes), are important for human physiology, which is often emphasized in diseased states caused by dysfunctional channels. For instance, K_V_7.1 is predominantly expressed in cardiomyocytes, and mutations to this channel is a known risk factor for developing cardiac arrhythmia (Barhanin et al., 1996, Sanguinetti et al., 1996, Tester and Ackerman, 2014, Brewer et al., 2020). K_V_7.2 and K_V_7.3 are broadly expressed in neurons where they form heterotetrameric K_V_7.2/7.3 channels, and mutations to these channels may give rise to epilepsy or chronic pain (Biervert et al., 1998, Wang et al., 1998, Nappi et al., 2020). K_V_7.4 is of particular importance in the auditory system where it forms homotetrameric channels responsible for a K^+^ conductance at the resting membrane potential of cochlear outer hair cells (OHCs) (Kubisch et al., 1999, Kharkovets et al., 2000, Rim et al., 2021). Mutations that perturb the trafficking or function of K_V_7.4 in OHCs are associated with a subtype of progressive hearing loss known as DFNA2 (Kubisch et al., 1999, Kharkovets et al., 2006, Gao et al., 2013, Rim et al., 2021). K_V_7.5 is more widely spread through the central nervous system and is particularly important in regulating excitability in the hippocampus (Schroeder et al., 2000, Tzingounis et al., 2010, Fidzinski et al., 2015). K_V_7.1, K_V_7.4 and K_V_7.5 are expressed in various smooth muscle cells such as vascular smooth muscle cells (VSMC), suggesting that several K_V_7 subtypes, including heterotetrameric K_V_7.4/7.5 channels, may contribute to the hyperpolarizing K^+^ current in VSMCs that promotes vasodilation by preventing Ca^2+^-dependent contraction (Ng et al., 2011, Mani et al., 2013, Stott et al., 2014, Bercea et al., 2021), The modulation of K_V_7.1 and K_V_7.2/7.3 channels by endogenous and pharmacological compounds has been extensively studied (Miceli et al., 2018, Wu and Larsson, 2020). However, less is known about the modulation of K_V_7.4 and K_V_7.5. Here we studied the effects of a class of channel modulators, polyunsaturated fatty acids (PUFAs), on K_V_7.4 and K_V_7.5, as these channels are expressed in excitable tissues and could be valuable pharmacological targets in different diseases.

There is emerging evidence that suggests that PUFAs influence the physiology of tissues that express K_V_7.4 and K_V_7.5 channels. For instance, a number of studies have found an inverse relationship between hearing loss and the PUFA plasma concentration, suggesting that the risk of impaired hearing decreases with an increased dietary intake of ω-3 PUFAs such docosahexaenoic acid (DHA) and eicosapentaenoic acid (EPA) (Dullemeijer et al., 2010, Gopinath et al., 2010, Curhan et al., 2014). Meta-analyses of randomized control trials have found that a multitude of beneficial cardiovascular outcomes, including anti-inflammatory and hypotensive effects, are associated with an increased intake of PUFAs (Miller et al., 2014, AbuMweis et al., 2018). There are likely several mechanisms that contribute to these PUFA effects. For instance, the protective PUFA effect on hearing loss has been attributed to cerebrovascular effects, reasoning that PUFAs improve circulation to the cochlea (Dullemeijer et al., 2010, Gopinath et al., 2010, Curhan et al., 2014). PUFA-induced vasodilation has in part been attributed to the activation of Ca^2+^-dependent and ATP sensitive K^+^ channels by PUFAs (Hoshi et al., 2013, Limbu et al., 2018, Bercea et al., 2021). However, little is known about the putative contribution of K_V_7.4 and K_V_7.5 channels to PUFA effects. We find this open question interesting because PUFAs have been shown to activate both K_V_7.1 and K_V_7.2/7.3 channels (Liin et al., 2015, Liin et al., 2016, Taylor and Sanders, 2017, Bohannon et al., 2019, Larsson et al., 2020). This PUFA-induced activation is mediated through a lipoelectric mechanism in which the PUFA tail inserts into the outer leaflet of the lipid bilayer adjacent to the channel, whereupon the negatively charged carboxyl head group of the PUFA interacts electrostatically with positively charged arginines in the voltage-sensing domain (VSD) of the channel (Liin et al., 2018, Yazdi et al., 2021). This electrostatic interaction facilitates the outward S4 movement causing a shifted voltage dependence of channel opening towards more negative voltages. However, the effect of PUFAs on K_V_7.4 and K_V_7.5 remains unstudied. In this study, we therefore aimed to characterize the response of K_V_7.4 and K_V_7.5 channels to PUFAs, in order to expand our understanding of how the K_V_7 family of channels responds to such lipids.

We report that PUFAs activate K_V_7.5 by shifting the voltage dependence of channel opening towards more negative voltages. Surprisingly, we find that PUFAs inhibit the K_V_7.4 channel by shifting the voltage dependence of channel opening towards more positive voltages. Thus, the K_V_7.5 channel’s response to PUFAs is largely in line with the responses that have previously been observed in K_V_7.1 and K_V_7.2/7.3, whereas the K_V_7.4 channel response is not. Our study expands our understanding of how members of the K_V_7 family respond to PUFAs and reveal subtype specific responses to these lipids.

## Results

### The PUFA docosahexaenoic acid activates hK_V_7.5, but inhibits hK_V_7.4

We began with investigating the effects of the physiologically abundant (Kim et al., 2014) PUFA DHA (molecular structure shown in Fig. 1A) on homotetrameric hK_V_7.5 or hK_V_7.4 channels expressed in *Xenopus* oocytes. The hK_V_7.5 channel was activated by 70 μM DHA, as was evident by the significant shift in the midpoint of the voltage dependence of channel opening (V_50_) towards more negative voltages (Fig. 1B, average shift of -21.5 ± 1.9 mV, *P* = < 0.0001). This allows hK_V_7.5 channels to open and conduct a K^+^ current at more negative voltages in the presence of DHA. 70 μM DHA did not cause a consistent change in the maximum conductance (G_max_) of the hK_V_7.5 channel (average relative ΔG_max_ was 1.06 ± 0.11, *P* = 0.59).

**Figure 1.**
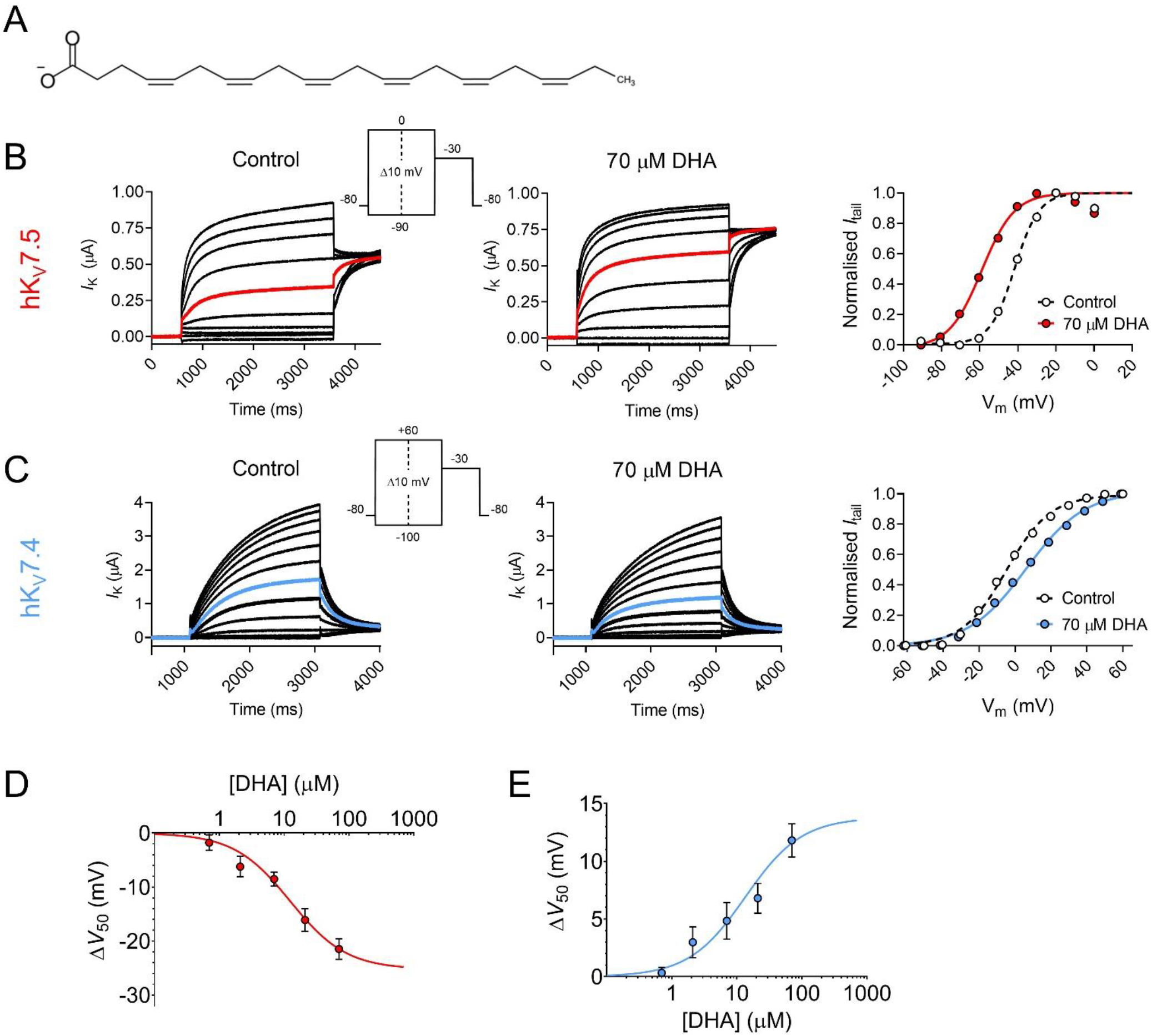
Docosahexaenoic acid activates hK_V_7.5 but inhibits hK_V_7.4. A) Molecular structure of DHA. B) Representative current family with corresponding G(V) curve of hK_V_7.5 in the absence (left) and presence (middle) of 70 µM DHA. Currents generated by the voltage protocol shown as inset. Red traces denote current generated by a test voltage to -40 mV. The G(V) curves (right) have been normalized between 0 and 1, as described in Materials and Methods, to better visualize shifts in V_50_. Curves represent Boltzmann fits (see Materials and Methods for details). V_50_ for this specific cell: V_50,ctrl_ = -41.7 mV, V_50,DHA_ = -59 mV. C) Same as in B but for hK_V_7.4. Blue traces denote current generated by a test voltage to 0 mV. V_50_ for this specific cell: V_50,ctrl_ = -4.8 mV, V_50,DHA_ = +6.2 mV. D-E) Concentration-response curve of the DHA effect on V_50_ of hK_V_7.5 (D) and hK_V_7.4 (E). Curves represent concentration-response fits (see Materials and Methods for details). Best fits: ΔV_50,max_ is -25.3 mV for hK_V_7.5 and +13.8 mV for hK_V_7.4. EC_50_ is 12 µM for hK_V_7.5 and 14 µM for hK_V_7.4. Data shown as mean ± SEM. n = 3-15. See also Figure 1 – Figure Supplement 1.

In clear contrast to the activating effect observed for hK_V_7.5, the hK_V_7.4 channel was inhibited by 70 μM DHA, as was seen by the significant shift in the V_50_ towards more positive voltages (Fig. 1C, average shift of +11.8 ± 1.4 mV, *P* = <0.0001). Furthermore, the application of DHA led to a more shallow slope of the G(V) curve (average slope factors were 13.1 ± 0.2 mV and 17.6 ± 0.5 mV in the absence and presence of 70 μM DHA, respectively). 70 μM DHA did not cause a consistent change in G_max_ of the hK_V_7.4 channel (average relative ΔG_max_ was 1.09 ± 0.08, *P* = 0.28). Because 70 μM DHA induced significant shifts in V_50_ of both hK_V_7.5 and hK_V_7.4, although in opposite directions, without affecting G_max_ we will throughout the remainder of this study focus our analysis of PUFA effects on V_50_, and G(V) curves reporting on PUFA effects will be normalized to facilitate visualization of V_50_ shifts.

The DHA-evoked shift in the V_50_ of hK_V_7.5 was significant at concentrations as low as 7 μM (Fig. 1D, -8.5 ± 1.3 mV, *P* = 0.0002). The concentration-response curve for the DHA effect on hK_V_7.5 predicts a maximal shift of -25.3 mV, with 12 μM required for 50% of the maximal shift (EC_50_ = 12 μM). The DHA effect on the V_50_ of hK_V_7.4 was also significant at 7 μM (Fig. 1E, +4.8 ± 1.6 mV, *P* = 0.01). The concentration-response curve for the DHA effect on hK_V_7.4 predicts a maximal shift of +13.8 mV, with an EC_50_ of 14 μM. The onset of the DHA effect was relatively fast for both the hK_V_7.5 and hK_V_7.4 channels, reaching a stable level within about 6 minutes (Figure 1 – Figure Supplement 1). The DHA effect proved difficult to wash out or reverse as re-perfusion with control solution or control solution supplemented with 100 mg/mL bovine serum albumin (BSA) only partially restored baseline current amplitude for hK_V_7.5 and hK_V_7.4 (Figure 1 – Figure Supplement 1). Altogether, the PUFA DHA has a concentration-dependent *activating* effect on the hK_V_7.5 channel, and a concentration-dependent *inhibitory* effect on the hK_V_7.4 channel.

### The hK_V_7.5 response, but not the hK_V_7.4 response, changes direction in an electrostatic manner

To further understand the molecular basis of the DHA response, we examined the importance of the charge of the DHA head group. Several previous studies identify that electrostatic interactions between positively charged residues in the VSD and a negatively charged head group on PUFAs (and their analogues) are fundamental for facilitating the activation of hK_V_7.1, hK_V_7.2/7.3 and some other K_V_ channels (Börjesson and Elinder, 2011, Liin et al., 2015, Liin et al., 2016, Liin et al., 2018, Martín et al., 2021). This is seen as a shift in V_50_ towards more negative voltages. The electrostatic PUFA effect on V_50_ can be tuned, from inducing negative to positive shifts in V_50_, by altering the charge of the PUFA head group (Börjesson et al., 2010, Liin et al., 2015). To investigate if the same electrostatic mechanism is at play in the responses of hK_V_7.5 and hK_V_7.4, we compared the DHA response of the channels with: 1) a DHA analogue with an uncharged methyl ester head group (DHA-me), and 2) a DHA analogue with a positively charged amine head group (DHA+).

70 μM of DHA-me did not significantly shift V_50_ of hK_V_7.5 (Fig. 2A, V_50_ was -0.2 ± 0.5 mV, *P* = 0.67). In contrast, 70 μM DHA+ brought on a small, but significant positive shift in V_50_ of hK_V_7.5 (Fig. 2A, ΔV_50_ was +4.5 ± 0.9 mV, *P* = 0.001). Thus, for hK_V_7.5 the negatively charged DHA facilitates activation, the uncharged DHA-me has no effect, and the positively charged DHA+ inhibits channel activation (effects summarized in Fig. 2B). This is in line with the lipoelectric mechanism that has been proposed to explain PUFA effects on the hK_V_7.1 and hK_V_7.2/7.3 channels (Liin et al., 2015, Liin et al., 2016).

**Figure 2.**
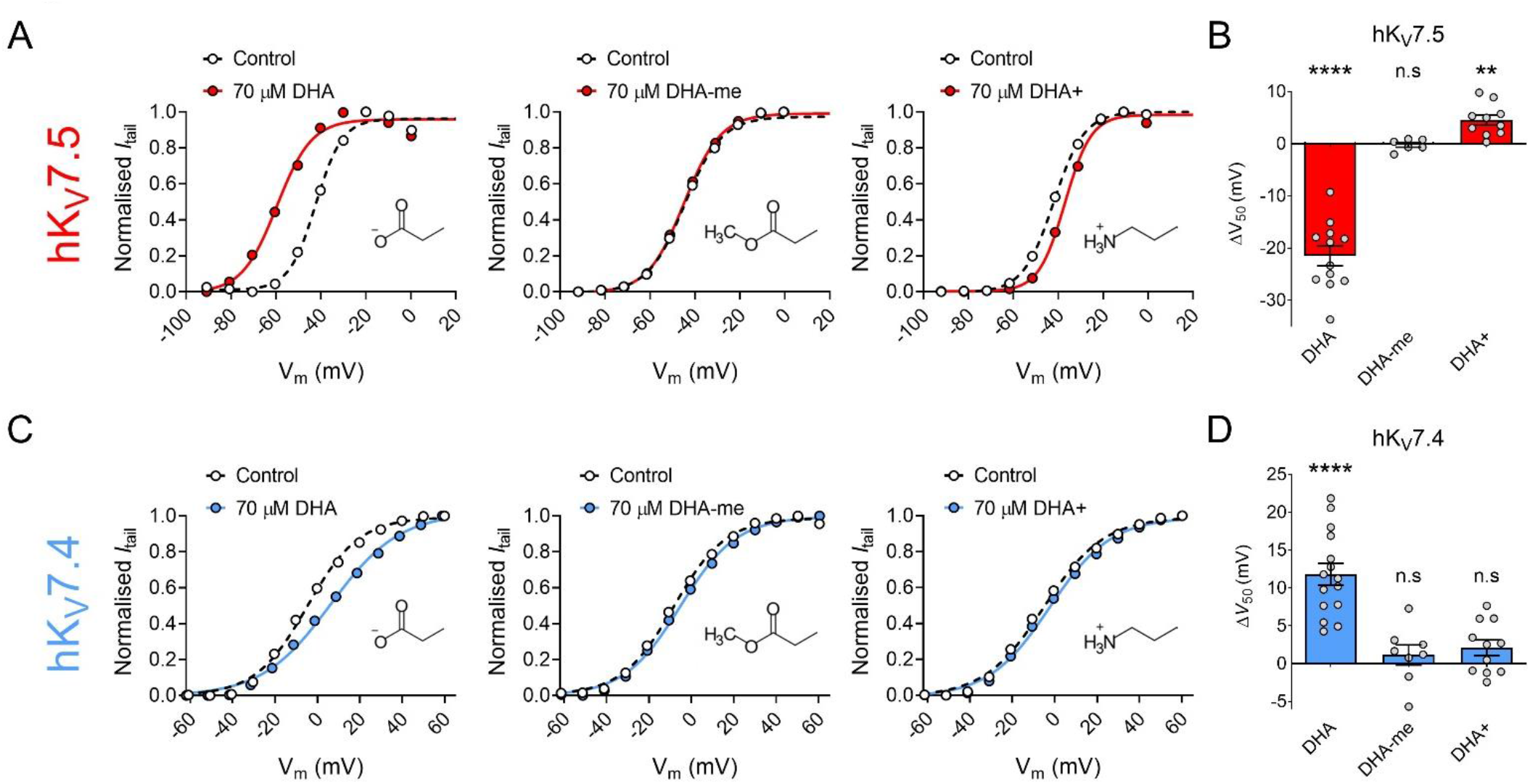
The impact of PUFA head group charge for hK_V_7.5 and hK_V_7.4 effects. A-B) Impact of the head group charge on the ability of DHA to shift V_50_ of hK_V_7.5, assessed by comparing the effect of negatively charged DHA, uncharged DHA-me, and positively charged DHA+ at 70 µM. A) Representative examples of the negative shift in V_50_ induced by DHA (left, same data as shown in Fig. 1B), the lack of effect of DHA-me (middle, V_50,ctrl_ = -44.6 mV; V_50,DHA-me_ = -44.9 mV), and the positive shift in V_50_ induced by DHA+ (right, V_50,ctrl_ = - 42 mV; V_50,DHA+_ = -37 mV). Molecular structure of head groups are shown as insets. B) Summary of responses to indicated DHA compound. Data shown as mean ± SEM. n = 6-12. Statistics denote one sample *t* test against a hypothetical mean of 0 mV. n.s denotes not significant, ** denotes *P* ≤ 0.01 **** denotes *P* ≤ 0.0001. C-D) same as in A-B but for hK_V_7.4. C) Representative examples of the depolarizing shift in V_50_ induced by DHA (left, same data as shown in Fig. 1C), the lack of effect of DHA-me (middle, V_50,ctrl_ = -8.3 mV; V_50,DHA-me_ = -5.2 mV), and the lack of effect of DHA+ (right, V_50,ctrl_ = -5 mV; V_50,DHA+_ = -2.3 mV). D) Summary of responses to indicated DHA compound. Data shown as mean ± SEM. n = 8-15. Statistics denote one sample *t* test against a hypothetical mean of 0 mV. n.s denotes not significant, **** denotes *P* ≤ 0.0001.

Neither 70 μM of DHA-me nor 70 μM of DHA+ had significant effects on V_50_ of hK_V_7.4 (Fig. 2C, ΔV_50_ was +1.1 ± 1.3 mV, *P* = 0.42 for DHA-me and +2.1 ± 1.0 mV, *P* = 0.068 for DHA+). Thus, while the negatively charged DHA caused a positive shift in V_50_ of hK_V_7.4, neither the uncharged DHA-me nor the positively charged DHA+ altered the V_50_ of hK_V_7.4 (effects summarized in bar graph of Fig. 2D). Even though a negative charge of the head group seems to be important for the effect, the responses are not in line with the lipoelectric mechanism, making hK_V_7.4 unique among the hK_V_7 channels.

### Both hK_V_7.5 and hK_V_7.4 respond broadly to PUFAs, although with varied magnitudes of responses

Next, we studied if the effects on hK_V_7.5 and hK_V_7.4 are specific to DHA or if they are shared among different PUFAs by testing a series of PUFAs, listed in Table I. These PUFAs vary in molecular properties such as the length of the tail and the number and position of double bonds in the tail. Figure 3A shows a plot of the magnitude of ΔV_50_ following exposure of hK_V_7.5 to PUFAs (at 70 μM) against the number of carbon atoms in each respective PUFA tail. All PUFAs, regardless of length (14-22 carbon atoms) significantly shifted V_50_ of hK_V_7.5 towards more negative voltages, although to different extents. The magnitude of the PUFA-induced ΔV_50_ of hK_V_7.5 followed a pattern of TTA < LA = AA < HTA ≤ EPA < DHA. One observation is that the ω-6 PUFAs LA and AA induced smaller shifts than the ω-3 PUFAs HTA, EPA and DHA. Both AA and EPA are 20 carbon atoms long. However, while AA is an ω-6 PUFA, EPA is an ω-3 PUFA (Table I). Figure 3B shows representative G(V) curves for hK_V_7.5 that highlight the larger shift in V_50_ evoked by 70 μM of EPA (left) than by 70 μM of AA (right). On average, EPA induced a shift of -17.9 ± 2.4 mV, which was significantly larger than the shift induced by AA (ΔV_50_ = -9.3 ± 1.7 mV; Fig. 3C; *P* = 0.015). These results suggest that in PUFAs of equal length, the ω-number has an impact on the magnitude of the PUFA response. We also compared the positively charged amine analogue DHA+ to the amine analogue of AA (AA+). The positive shift in V_50_ of hK_V_7.5 by 70 μM AA+ was slightly smaller than that observed with DHA+, and did not differ significantly from a hypothetical shift of 0 mV (Fig. 3C, AA+ ΔV_50_ = +1.9 ± 1.4 mV, *P* = 0.2). However, there was also no statistical difference in the effect induced by DHA+ and AA+ (*P* = 0.13), which indicates that the importance of the ω-number for amine analogues should be interpreted with caution. Altogether, these experiments show that many PUFAs activate the hK_V_7.5 channel and suggest that the hK_V_7.5 channel shows a preference towards ω-3 over ω-6 PUFAs.

**Table I.**
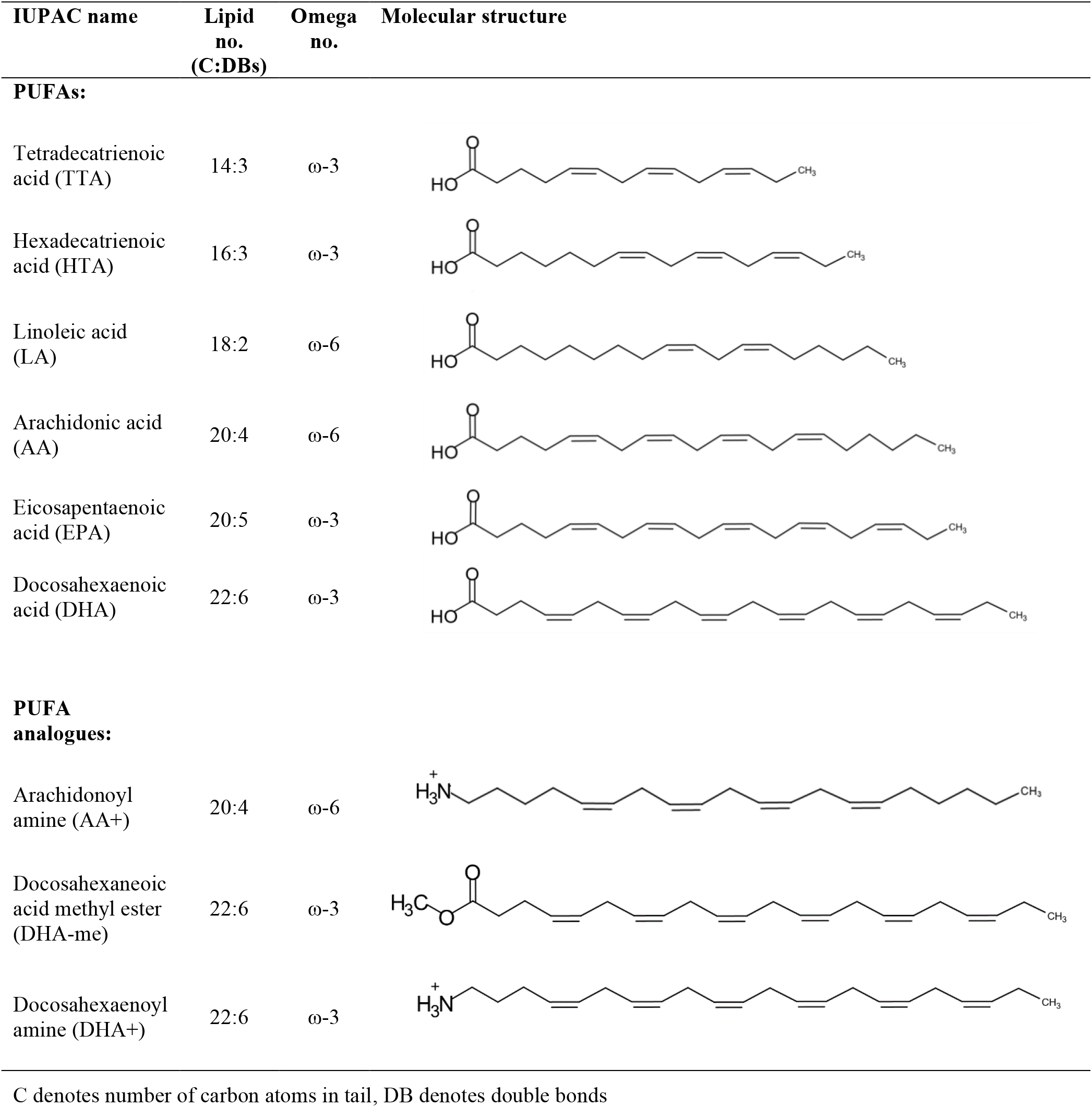
List of PUFAs and PUFA analogues used in this study.

| IUPAC name | Lipid no.<br>(C:DBs) | Omega no. | Molecular structure |
| --- | --- | --- | --- |
| **PUFAs:** |  |  |  |
| Tetradecatrienoic acid (TTA) | 14:3 | ω-3 | |
| Hexadecatrienoic acid (HTA) | 16:3 | ω-3 | |
| Linoleic acid (LA) | 18:2 | ω-6 | |
| Arachidonic acid (AA) | 20:4 | ω-6 | |
| Eicosapentaenoic acid (EPA) | 20:5 | ω-3 | |
| Docosahexaenoic acid (DHA) | 22:6 | ω-3 | |
| **PUFA analogues:** |  |  |  |
| Arachidonoyl amine (AA+) | 20:4 | ω-6 | |
| Docosahexaneic acid methyl ester (DHA-me) | 22:6 | ω-3 | |
| Docosahexaenoyl amine (DHA+) | 22:6 | ω-3 | |

C denotes number of carbon atoms in tail, DB denotes double bonds

**Figure 3.**
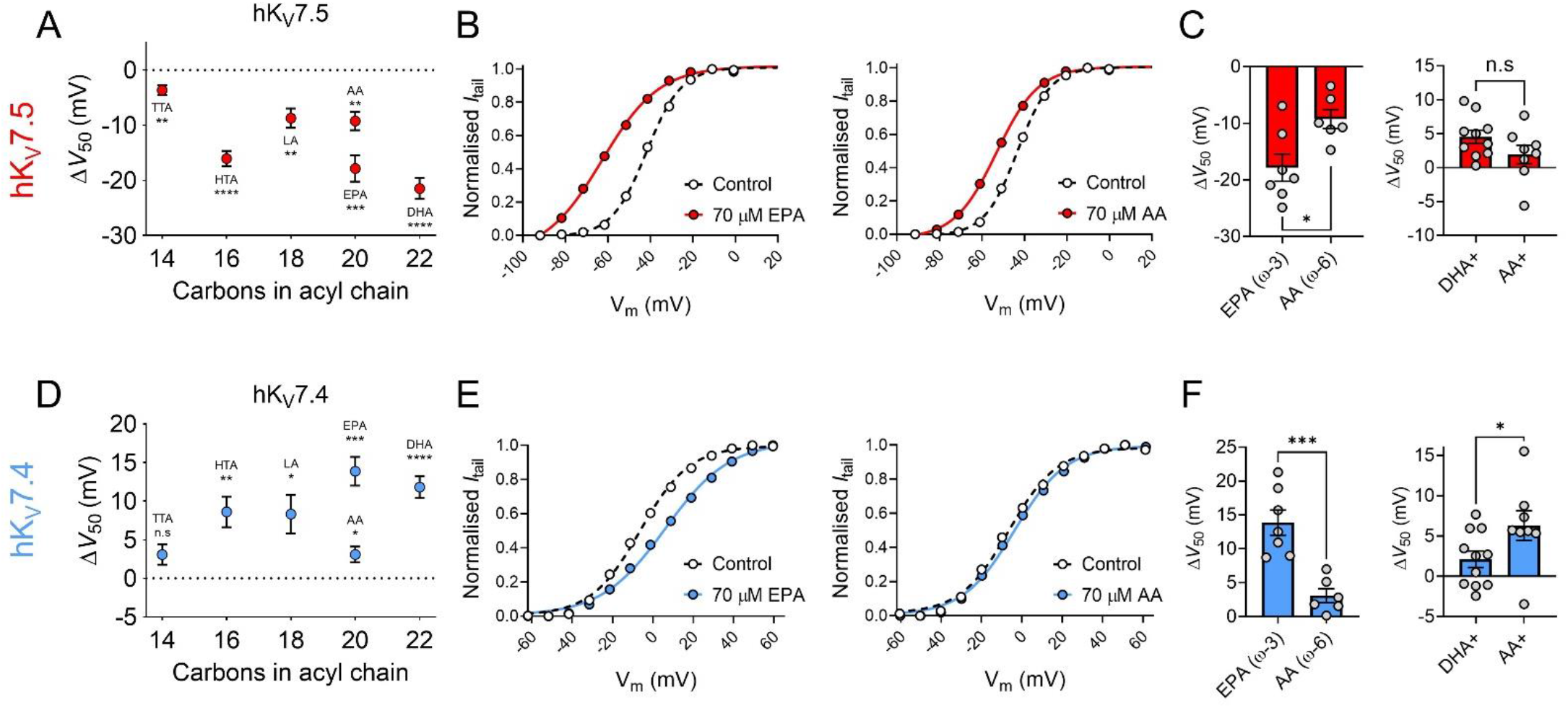
The impact of PUFA tail properties for hK_V_7.5 and hK_V_7.4 effects. A-C) Impact of PUFA tail properties for the ability of PUFAs to shift V_50_ of hK_V_7.5, assessed by comparing the response to 70 µM of indicated PUFAs (see Table I for molecular structures). A) Average effect of indicated PUFAs. Data shown as mean ± SEM. n = 6-8. Statistics denote one sample *t* test against a hypothetical mean of 0 mV n.s denotes not significant, ** denotes *P* ≤ 0.01, *** denotes *P* ≤ 0.001, **** denotes *P* ≤ 0.0001. B) Representative examples of response to 70 µM of the ω-3 PUFA EPA (left, V_50,ctrl_ = -41.9 mV; V_50,EPA_ = -62.8 mV) and ω-6 PUFA AA (middle, V_50,ctrl_ = -43.3 mV; V_50,AA_ = -53.1 mV). Bar graphs with comparison of average response to 70 µM of EPA and AA (n = 6-7) and DHA+ and AA+ (n = 8-10). Statistics denote Student’s *t* test. n.s denotes not significant, * denotes *P* ≤ 0.05. D-F) same as in A-C but for hK_V_7.4. D) n = 6-8. E) Left: V_50,ctrl_ = -6.3 mV; V_50,EPA_ = +6.2 mV. Middle: V_50,ctrl_ = -6.5 mV; V_50,AA_ = -3.6 mV. F) n = 6-7 and n = 8-11, respectively. * denotes *P* ≤ 0.05, ** denotes *P* ≤ 0.01, *** denotes *P* ≤ 0.001, **** denotes *P* ≤ 0.0001.

Figure 3D shows a plot of the magnitude of ΔV_50_ following exposure to PUFAs (at 70 μM) against the number of carbon atoms in each respective PUFA tail for the hK_V_7.4 channel. The shortest PUFA TTA did not evoke a significant shift in V_50_ (ΔV_50_ = +3.1 ± 1.3 mV, *P* = 0.051). All remaining PUFAs (at a concentration of 70 μM), however, significantly shifted V_50_ of hK_V_7.4 towards positive voltages. The magnitude of the PUFA-induced ΔV_50_ of hK_V_7.4 followed a pattern of TTA < AA < LA ≤ HTA < DHA < EPA with no clear pattern in magnitude of effect based on the ω-number of the PUFA tail. Although the ω-3 PUFA EPA induced a larger shift than the ω-6 PUFA AA (Fig. 3E-F, EPA ΔV_50_ = +13.9 ± 1.9 mV; AA ΔV_50_ = +3.1 ± 1.0 mV, *P* = < 0.001), AA+ caused a larger shift in V_50_ compared to DHA+ (Fig. 3F, DHA+ ΔV_50_ = +2.1 ± 1.0 mV; AA+ ΔV_50_ = +6.3 ± 1.8 mV, *P* = 0.049). Moreover, the ω-6 PUFA LA induced comparable shifts in V_50_ to those of the ω-3 PUFA HTA. Altogether, these experiments show that many PUFAs inhibit the hK_V_7.4 channel, with no obvious pattern of which PUFAs hK_V_7.4 shows a preference towards.

### The DHA response of hK_V_7.5 is greater than that of other hK_V_7 subtypes

We have in previous studies shown that 70 μM of DHA shifts V_50_ of the hK_V_7.1 and hK_V_7.2/7.3 channels by about -9 mV (Fig. 4A; hK_V_7.1 ΔV_50_ = -9.3 ± 0.9 mV as reported in (Liin et al., 2015); hK_V_7.2/7.3 ΔV_50_ = -9.3 ± 1.7 mV as reported in (Liin et al., 2016)). Thus, the extent of the hK_V_7.5 shift induced by 70 μM DHA is greater than that of hK_V_7.1 or hK_V_7.2/7.3 (Fig. 4A). Our experiments indicated the necessity of a negatively charged DHA head group to elicit the activating effect on hK_V_7.5 (see Fig. 2A). A negatively charged head group is promoted by alkaline pH, which triggers proton dissociation (Hamilton, 1998, Börjesson et al., 2008). In a previous study, the apparent pKa of DHA near hK_V_7.1 (i.e. the pH at which 50% of the maximal DHA effect is seen, interpreted as the pH at which 50% of the DHA molecules in a lipid environment are negatively charged) was determined to be pH 7.7 (Liin et al., 2015). One possible underlying cause of the larger DHA effect on hK_V_7.5 is that the local pH at hK_V_7.5 promotes DHA deprotonation, thus rendering a greater fraction of the DHA molecules negatively charged and capable of activating the channel. To test this, we assessed the effect of 70 μM DHA on hK_V_7.5 with the extracellular pH adjusted to either more alkaline (pH = 8.2) or acidic (pH = 6.5, or 7.0) values. At pH 8.2, at which a majority of DHA molecules are expected to be deprotonated, DHA shifted V_50_ almost two-fold greater than at physiological pH (pH 7.4 ΔV_50_ = -21.5 ± 1.9 mV; pH 8.2 ΔV_50_ = -44.4 ± 3.2 mV, *P* < 0.0001, Student’s *t* test, representative example in Fig. 4B). The magnitude of the DHA effect was reduced as the pH was gradually titrated towards acidic pH (Fig. 4C), with no shift in V_50_ by 70 μM DHA at pH 6.5 (ΔV_50_ = +0.1 ± 1.0 mV, *P* = 0.96). The pH dependence of ΔV_50_ induced by DHA for the hK_V_7.1 channel is also plotted in Figure 4C, to allow for comparison between the two hK_V_7 subtypes. While there is a clear difference in the extrapolated maximum shifts for the two channels (hK_V_7.1 ΔV_50,max_ = -26.3 mV with 95% CI [-18.4, - 34.3]; hK_V_7.5 ΔV_50,max_ = -57.9 mV with 95% CI [-47.6, -68.2]), the pH required to induce 50% of the maximum ΔV_50_ (i.e, the apparent pKa values) are similar (inset of Fig. 4C, hK_V_7.1 apparent pKa = 7.68 with 95% CI [7.26, 7.89]; hK_V_7.5 apparent pKa = 7.67 with 95% CI [7.52, 7.90]). This indicates that the difference in the DHA effect between hK_V_7.1 and hK_V_7.5 is a question of increased magnitude rather than a matter of different local pH, as we would expect the DHA effect of the hK_V_7.5 channel to exhibit a lower apparent pKa if the channel was promoting DHA deprotonation.

**Figure 4.**
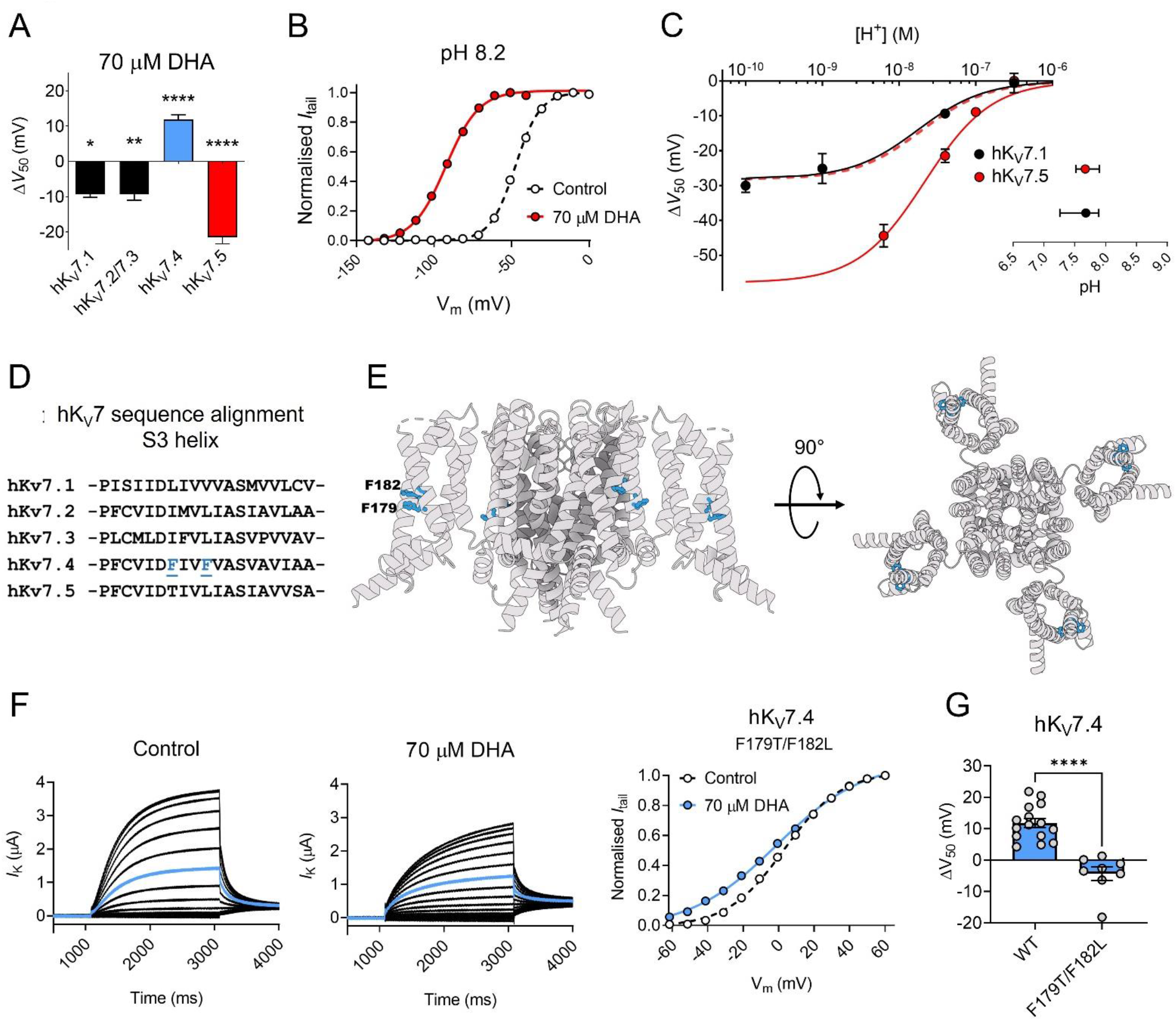
Extracellular pH and S3 residues tune the DHA response of hK_V_7.5 and hK_V_7.4, respectively. A) Comparison of the V_50_ shift of hK_V_7 subtypes in response to 70 µM DHA. Data for hK_V_7.1 and hK_V_7.2/7.3 shown as reported in (Liin et al., 2015, Liin et al., 2016). Data for hK_V_7.4 and hK_V_7.5 from the present study. Data shown as mean ± SEM. n = 3-15. Statistics denote one sample *t* test against a hypothetical mean of 0 mV. * denotes *P* ≤ 0.05, ** denotes *P* ≤ 0.01, **** denotes *P* ≤ 0.0001. B-C) Altering extracellular pH tunes the shift in V_50_ of hK_V_7.5 induced by 70 µM DHA, with a greater response observed at more alkaline pH. B) Representative example of the DHA response at pH 8.2 (V_50,ctrl_ = -47.6 mV; V_50,DHA_ = -92.4 mV). C) pH-response curve for the DHA effect on V_50_. Data shown as mean ± SEM. n = 3-9. Curves represent pH-response fits (see Materials and Methods for details). Best fit for hK_V_7.5: ΔV_50,max_ is -57.9 mV. Data for hK_V_7.1 as reported in (Liin et al., 2015) is included for comparison (Best fit for hK_V_7.1: ΔV_50,max_ is -26.3 mV.) Dashed line denotes the pH-response curve for hK_V_7.5 normalized to ΔV_50,max_ for hK_V_7.1 to illustrate the comparable apparent pKa. Inset shows apparent pKa with 95% CI: hK_V_7.1 apparent pKa = 7.68 with 95% CI [7.26, 7.89]; hK_V_7.5 apparent pKa = 7.67 with 95% CI [7.52, 7.90]. Note that inconsistent behavior of hK_V_7.5 under control conditions at pH higher than 8.2 prevented us from determining the DHA effect on hK_V_7.5 at pH 9 and 10. D) Sequence alignment of the S3 helix of hK_V_7 subtypes, identifying two phenylalanine residues, F179 and F182, unique to hK_V_7.4. E) Structural model (PDB: 7BYL) of hK_V_7.4 with position of F179 and F182 marked. Left, side view, right top view. F-G) Substitution of F179 and F182 in hK_V_7.4 to the hK_V_7.5 counterparts (F179T/F182L) alters the response to 70 µM of DHA. F) Representative current family with corresponding G(V) curve of hK_V_7.4_F179/F182L in the absence (left) and presence (middle) of 70 µM DHA. Blue traces denote current generated by a test voltage to 0 mV. Curves in G(V) plots (right) represent Boltzmann fits. V_50_ for this specific cell: V_50,ctrl_ = +3.2 mV, V_50,DHA_ = -1.1 mV. G) Summary of average response to 70 µM DHA on indicated constructs. Data shown as mean ± SEM. n = 8-15. Statistics denote Student’s *t* test. **** denotes *P* ≤ 0.0001.

### Phenylalanine residues in the S3 helix of hK_V_7.4 contribute to the PUFA response

We next turned our attention towards structural components of the hK_V_7.4 channel that may contribute to its unusual PUFA response. Given that the ΔV_50_ effect of PUFA on hK_V_7.1 has been shown to be mediated through PUFA-gating charge interactions at the VSD (Liin et al., 2018, Yazdi et al., 2021), we compared amino acid composition in the VSD between hK_V_7 channels. Sequence alignment of the S3 helix of hK_V_7.1-7.5 revealed two phenylalanine residues (F179 and F182) exclusive to hK_V_7.4 (Fig. 4D-E). We hypothesized that the presence of these two residues with bulky, aromatic side chains may in some way be responsible for the unusual response of hK_V_7.4 to PUFA. By means of site-directed mutagenesis, we substituted these phenylalanine residues for their hK_V_7.5 counterparts, a threonine and a leucine, creating the hK_V_7.4 F179T/F182L channel. The hK_V_7.4 F179T/F182L mutant behaved fairly similar to the wild-type hK_V_7.4 channel under control conditions, with a V_50_ shifted about 10 mV towards more positive voltages than for the wild-type channel (Table II). We then probed the response of the F179T/F182L mutant channel to 70 µM DHA. Representative K^+^ currents through hK_V_7.4 F179T/F182L before and after exposure to 70 μM DHA, as well as the corresponding G(V) curves, are shown in Figure 4F. The positive shift in V_50_ normally seen in wild-type hK_V_7.4 by 70 μM DHA was absent in hK_V_7.4 F179T/F182L (Fig. 4F-G, ΔV_50_ = -4.3 ± 2.2 mV, *P* = 0.086). Thus, while the F179T/F182L mutation did not endow the hK_V_7.4 channel with a substantial activating response to DHA typical of other hK_V_7 channels, it was sufficient to abrogate the inhibitory response to DHA observed in the wild-type hK_V_7.4 channel (*P* < 0.0001, Student’s *t* test).

**Table II.** Biophysical properties of tested constructs under control conditions.

| Channel | Variant | $V_{50}$ (mV)<br>Mean ± SEM | Slope (mV)<br>Mean ± SEM | n |
| --- | --- | --- | --- | --- |
| hK<sub>v</sub>7.4 | Wild-type | -11.1 ± 1.4 | 13.1 ± 0.2 | 20 |
| | F179T/F182L | +0.5 ± 2.5 | 15.8 ± 0.5 | 8 |
| | F182L | -21.0 ± 2.9 | 12.0 ± 0.5 | 10 |
| | S273A | -15.9 ± 3.1 | 14.1 ± 1.9 | 15 |
| hK<sub>v</sub>7.5 | Wild-type | -42.2 ± 0.7 | 8.4 ± 0.2 | 23 |
| hK<sub>v</sub>7.4/7.5 | Wild-type | -22.9 ± 0.6 | 10.8 ± 0.2 | 18 |

n denotes number of cells. $V_{50}$ and slope were determined from Boltzmann fits, as described in Material and Methods.

### hK_V_7.4/7.5 co-expression and disease-associated hK_V_7.4 mutations influence the response to DHA

We next characterized the DHA response of oocytes co-injected with cRNAs encoding hK_V_7.4 and hK_V_7.5 (referred to as hK_V_7.4/7.5) to allow for the potential formation of heteromeric channels containing hK_V_7.4 and hK_V_7.5 subunits. Oocytes co-injected with hKv7.4 and hKv7.5 generated currents with biophysical properties that fell between those of each homomer (hK_V_7.4 V_50_ = -11.1 ± 1.4 mV; hK_V_7.5 V_50_ = -42.2 ± 0.7 mV; hK_V_7.4/7.5 V_50_ = -22.9 ± 0.6 mV, Table II). This is in agreement with previous studies showing a V_50_ of co-expressed hK_V_7.4/7.5 channels intermediate to that of homomeric channels (Brueggemann et al., 2011). In addition, 70 μM of DHA induced an effect on hK_V_7.4/7.5 intermediate to that of homomeric hK_V_7.4 and hK_V_7.5 channels, with no effect on V_50_ (Fig. 5A-B, G, ΔV_50_ = -0.4 ± 0.6 mV, *P* = 0.53).

**Figure 5.**
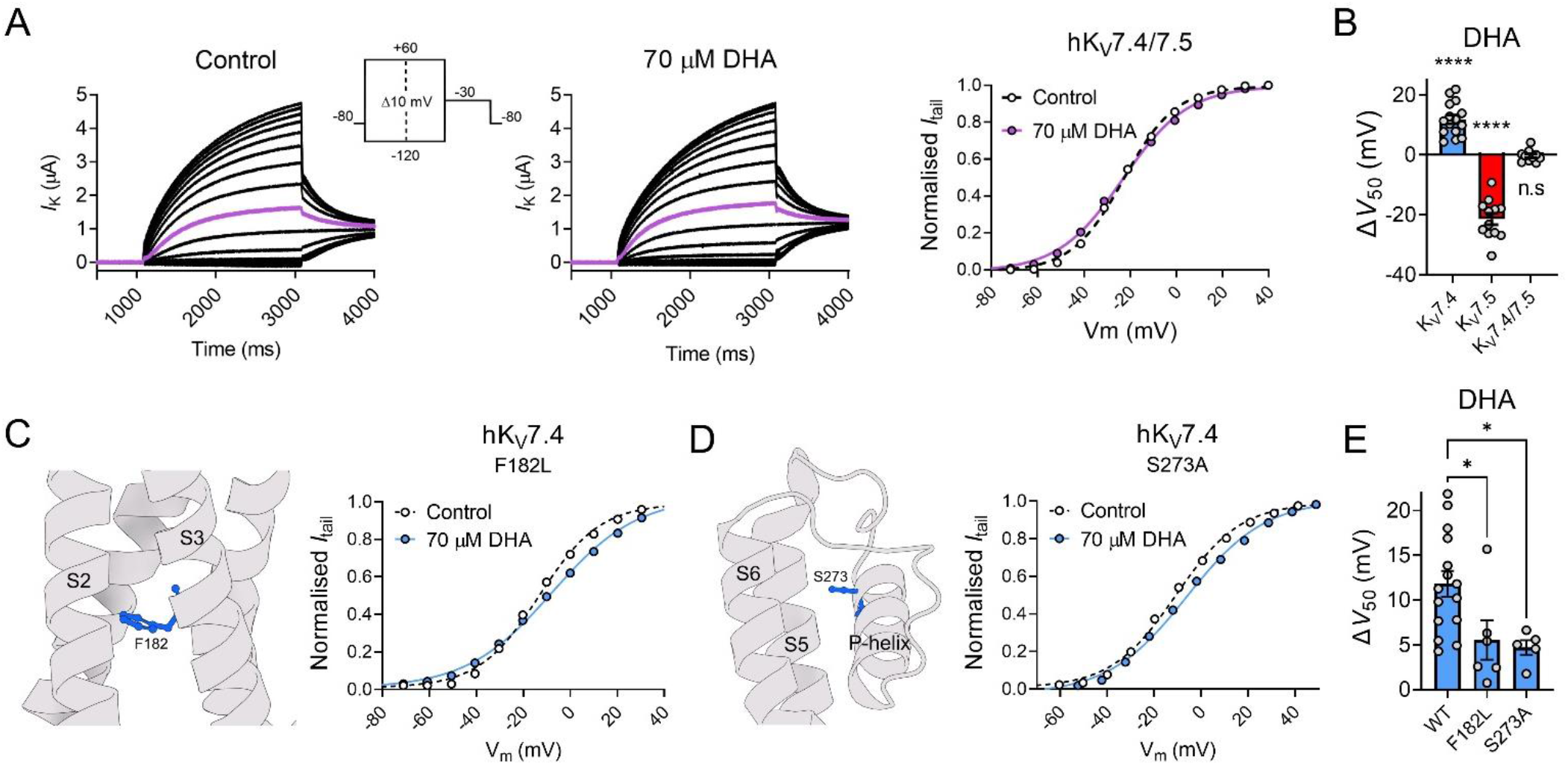
hK_V_7.4/7.5 co-expression and disease-associated hKv7.4 mutations influence the response to DHA. A) Representative current family with corresponding G(V) curve of hK_V_7.4/7.5 in the absence (left) and presence (middle) of 70 µM DHA. Purple traces denote current generated by a test voltage to -20 mV. Curves represent Boltzmann fits. V_50_ for this specific cell: V_50,ctrl_ = -22.8 mV; V_50,DHA_ = -23.1 mV. B) Summary of response of hK_V_7.4/7.5 to 70 µM DHA. Data shown as mean ± SEM. n = 10-15. Statistics denote one sample *t* test against a hypothetical mean of 0 mV. n.s denotes not significant, **** denotes *P* ≤ 0.0001. C-D) Impact of hK_V_7.4 mutations F182L (C) and S273A (D) on the response to DHA. Structural model (PDB: 7BYL) of hK_V_7.4 with position of F182 and S273 marked. Representative G(V) curve of indicated hK_V_7.4 mutants in the absence and presence of 70 µM DHA. Curves represent Boltzmann fits. For these specific cells: F182L: V_50,ctrl_ = -13.1 mV; V_50,DHA_ = -7.7 mV. For S273A: V_50,ctrl_ = -10.7 mV; V_50,DHA_ = -6.1 mV. E) Summary of response of WT hK_V_7.4 and indicated mutants to 70 µM DHA. Data shown as mean ± SEM. n = 5-15. Statistics denote one-way ANOVA followed by Dunnett’s multiple comparisons test to compare the response of mutants to that of the wild-type. * denotes *P* ≤ 0.05.

Several mutations in the KCNQ4 gene (encoding hK_V_7.4) have been identified in humans and are often linked to DFNA2 non-syndromic hearing loss (Jung et al., 2019, Rim et al., 2021), although the possible contribution to pathology remains to be determined for many mutations. Notably, the F182L and S273A missense mutations in the VSD of hK_V_7.4 (Fig. 5C-D), which are suspected to be linked to impaired hearing (Kim et al., 2011, Jung et al., 2019), exchanges the native hK_V_7.4 residue for the hK_V_7.5 counterpart. Under control conditions, we found both mutants to behave fairly similar to wild-type hK_V_7.4 (Table II). The S273A mutant had a V_50_ comparable to wild-type hK_V_7.4 whereas the V_50_ of the F182L mutant was shifted about 10 mV towards more negative voltages compared to wild-type hK_V_7.4 (Table II). However, both mutations impaired the hK_V_7.4 response to DHA. 70 µM of DHA shifted V_50_ of the F182L mutant by only +5.5 ± 2.2 mV and the S273A mutant by +4.7 ± 0.8 mV (Fig. 5C-E). Altogether, these experiments suggest that co-expression of hK_V_7.4 and hK_V_7.5 subunits, or substitution of specific residues in hK_V_7.4 to the hK_V_7.5 counterpart, alter the DHA response of hK_V_7.4 to approach that of hK_V_7.5.

## Discussion

This study finds that the activity of hK_V_7.4 and hK_V_7.5 channels are modulated by PUFAs. As such, our study expands upon the understanding of the modulation of the hK_V_7 family of ion channels by this class of lipids. Importantly, we find that both hK_V_7.4 and K_V_7.5 show surprising responses to PUFA, compared to previously reported effects on other hK_V_7 subtypes. hK_V_7.4 is the only hK_V_7 subtype for which PUFAs induce channel inhibition, whereas ω-3 PUFAs induce unexpectedly large hK_V_7.5 channel activation. Experiments on co-expressed hK_V_7.4/7.5 channels and hK_V_7.4 channels carrying hK_V_7.5 mimetic disease-associated mutations indicate responses to PUFAs that are intermediate of the hK_V_7.4 and hK_V_7.5 homomers. The diverse responses of hK_V_7 subtypes to PUFAs demonstrate the importance of carrying out functional experiments on each of the channel subtypes to allow for an understanding of subtype variability in channel modulation.

What mechanistic basis may underlie PUFA activation of hK_V_7.5? We find that the pattern of how hK_V_7.5 responds to PUFA conforms to the lipoelectric mechanism that has previously been described for the hK_V_7.1 and hK_V_7.2/7.3 channels, with a hyperpolarizing shift in V_50_ induced by negatively charged PUFAs, a depolarizing shift in V_50_ induced by a positively charged PUFA analogue, and no effect of an uncharged PUFA analogue. Therefore, we find it likely that PUFAs activate also hK_V_7.5 through electrostatic interactions with positively charged gating charges in the VSD. However, ω-6 PUFAs are less effective than ω-3 PUFAs at activating the hK_V_7.5 channel. This is different from the hK_V_7.1/KCNE1, for which ω-6 or ω-9 PUFAs were identified as the best activators of hK_V_7.1/KCNE1 (Bohannon et al., 2019). Although the role of the ω numbering in determining the extent of PUFA effects on hK_V_7.5 should be interpreted with caution (for instance because the PUFAs also vary in their number of double bonds, see Table I), it is clear that hK_V_7.5 and hK_V_7.1/KCNE1 do not follow the same response pattern to different PUFAs. Work on hK_V_7.1/KCNE1 has demonstrated that the KCNE1 subunit impairs the DHA response of hK_V_7.1 by altering the pH in the vicinity of the channel to promote DHA protonation (Liin et al., 2015, Larsson et al., 2018). This led us to test if the larger DHA response of the hK_V_7.5 channel was caused by a difference in the local pH compared to hK_V_7.1-7.3, albeit in a manner promoting DHA deprotonation. However, the similar apparent pKa values of DHA for hK_V_7.1 and hK_V_7.5 discards this hypothesis. Instead, we find a larger magnitude of the DHA effect on hK_V_7.5 at all pH values compared to Kv7.1. Interestingly, a residue at the top of S5 of K_V_7.1, Y278, which was recently identified as an important “anchor point” for binding the ω-6 PUFA LA near the VSD to allow for efficient shifts in V_50_ (Yazdi et al., 2021), is instead a phenylalanine (F282) in hK_V_7.5. A phenylalanine mutation of this “anchor point” in K_V_7.1 (Y278F) led to a drastic decrease in apparent affinity of LA for hK_V_7.1, possibly by removing hydrogen bonding between the head group of LA and the hydroxyl group of the tyrosine side chain (Yazdi et al., 2021). However, because the Y278F mutation of hK_V_7.1 also reduced the apparent affinity of DHA for hK_V_7.1 (Yazdi et al., 2021), and DHA was the most potent of the PUFAs we tested on hK_V_7.5 in this study, sequence variability at this position is not likely to underlie the relatively larger effect of DHA on K_V_7.5 compared to K_V_7.1. Thus, the nature of PUFA binding to hK_V_7 channels is more complex and there may be other stabilizing residues in the PUFA interaction site of the hK_V_7.5 channel.

We find that the hK_V_7.4 channel is inhibited by PUFAs, and that the direction of the effect is not reversed by reversing the charge of the PUFA head group. Therefore, the PUFA effect on hK_V_7.4 does not conform to the lipoelectric mechanism proposed for other Kv7 channels. Additionally, we did not find any relationship between PUFA tail properties and the response of hK_V_7.4 to PUFAs, other than that all PUFAs with significant effects shifted V_50_ towards more positive voltages. A cryo-EM structure for hK_V_7.4 was recently reported (PDB: 7BYL; (Li et al., 2021)), which reveals a major structural difference of putative importance to PUFA-hK_V_7 interactions when comparing the hK_V_7.4 structure to the hK_V_7.1 structure (PDB: 6UZZ; (Sun and MacKinnon, 2020)). The entire VSD of hK_V_7.4 is rotated 15° clockwise relative to the pore domain of the channel, which changes the geometry at a corresponding PUFA interaction site at the interface between the VSD and pore domain of hK_V_7.1 (Yazdi et al., 2021). One possibility is that the rotation of the VSD in hK_V_7.4 impairs PUFA interaction with this site, which could explain the lack of activating PUFA effects on hK_V_7.4. Another contributing factor to the unusual PUFA response of hK_V_7.4 may be the presence of bulky phenylalanines in S3 of hK_V_7.4, which directly or indirectly may alter PUFA interaction with the VSD. Moreover, PUFAs have been shown to inhibit other voltage-gated ion channels (Elinder and Liin, 2017, Bohannon et al., 2020). For instance, DHA has been suggested to inhibit K_V_1.1 and K_V_1.5 channels by interacting with hydrophobic residues in the channel pore (Decher et al., 2010, Bai et al., 2015). It is possible that a comparable interaction site for PUFAs, through which PUFAs mediate their inhibitory actions, exists in the hK_V_7.4 channel. Potential PUFA interaction sites underlying effects on hK_V_7.4 deserve further scrutiny in the future.

What potential physiological implications may our findings have? Increased consumption of foods rich in PUFAs have long been associated with improved cardiovascular health (Bang et al., 1980, Kagawa et al., 1982, Saravanan et al., 2010) and multiple mechanisms have been proposed including those mediated by the immune system, the endothelium and vascular smooth muscle tissues (Massaro et al., 2008). Several ion channels have been studied as the molecular correlates of PUFA-mediated vasodilatory mechanisms in both endothelial cells and VSMCs (Bercea et al., 2021). Among others, PUFA induced inhibition of L-type Ca_V_ channels (Engler et al., 2000) and PUFA induced activation of large conductance Ca^2+^-activated K^+^ (BK) channels contribute to the vasodilation induced by PUFA (Hoshi et al., 2013, Limbu et al., 2018). Intriguingly, a recent study by Limbu *et al*. (Limbu et al., 2018) observed an endothelium-independent residual relaxation of rodent arteries induced by ω-3 PUFAs despite pre-treatment with several channel inhibitors, suggesting there may be another mechanism underlying PUFA-mediated vasorelaxation that does not involve BK channels. Our finding that hK_V_7.5 is activated by PUFAs raises the possibility that hK_V_7.5 subunits contribute to the residual vasorelaxation observed by Limbu and colleagues.

Besides cardiovascular effects, an increased intake of ω-3 PUFAs has also indicated a potential protective role against hearing loss (Dullemeijer et al., 2010, Gopinath et al., 2010, Curhan et al., 2014). While these studies propose the protective mechanism of PUFAs on hearing stem from vascular effects that improve cochlear perfusion, no direct mechanism of PUFAs on the hearing organ was investigated. Based on our findings, it would be unlikely that the decreased risk of hearing loss is due to direct actions of PUFAs on hK_V_7.4 channels expressed in OHCs, given that we find the channel to be inhibited by PUFAs. However, PUFA-induced activation of K_V_7 channels (e.g. K_V_7.1 and K_V_7.5) in VSMC could possibly contribute to improved cochlear perfusion. Of note, the two DFNA2-associated missense mutations to hK_V_7.4 we examined, F182L and S273A, showed intrinsic biophysical behavior comparable to that of wild-type hK_V_7.4 (see Table II). This is in overall agreement with previous studies of these mutants performed in other expression systems showing voltage dependence of channel opening approximate to that of the wild-type channel (Kim et al., 2011, Jung et al., 2019). Jung and colleagues found that the S273A mutant reduces the average whole-cell current densities to less than 50% of the wild-type (Jung et al., 2019), which indicates that S273A may be a risk factor for development of hearing loss through its limited ability to generate K^+^ currents. However, whether F182L acts as a risk factor in DFNA2 hearing loss or should rather be considered a benign missense mutation (Kim et al., 2011) remains to be determined. We find that both the F182L and S273A mutant display impaired response to DHA, which, if anything, is expected to preserve channel function in the presence of PUFAs.

To conclude, we find that hK_V_7.4 and hK_V_7.5 respond in opposing manners to PUFAs. The hK_V_7.5 channel’s response is largely in line with the responses that have previously been observed in other hK_V_7 channels, whereas the hK_V_7.4 channel response is not. Altogether, our study expands our understanding of how the activity of different members of the hK_V_7 family are modulated by PUFAs and demonstrates different responses of different hK_V_7 subtypes. Further studies are needed to determine the mechanistic basis for the unusual hK_V_7.4 response and to evaluate putative physiological importance of the effects described in this study in more complex experimental systems.

## Materials and Methods

### Test compounds

All chemicals were purchased from Sigma-Aldrich, Stockholm, Sweden, unless stated otherwise. The PUFAs used in this study include: tetradecatrienoic acid (TTA, 14:3Δ5,8,11, Larodan, Stockholm, Sweden), hexadecatrienoic acid (HTA, 16:3Δ7,10,13, Larodan, Stockholm, Sweden), linoleic acid (LA, 22:6Δ4,7,10,13,16,19), arachidonic acid (AA, 20:4Δ5,8,11,14), eicosapentaenoic acid (EPA, 20:5Δ5,8,11,14,17) and docosahexaenoic acid (DHA, 22:6Δ4,7,10,13,16,19). PUFA analogues used in this study include: docosahexaenoic acid methyl ester (DHA-me), arachidonoyl amine (AA+), and docosahexaenoyl amine (DHA+). AA+ and DHA+ were synthesized *in house* as previously described (Börjesson et al., 2010). Table I summarizes the molecular properties of PUFAs and PUFA analogues used. PUFAs and PUFA analogues were solved in 99.9% EtOH and diluted to their final concentrations in extracellular recording solution. To prevent potential degradation of arachidonic acid, recording solution was supplemented with 5 mM of the cyclooxygenase inhibitor indomethacin. Previous experiments with radiolabeled fatty acids have found that up to 30% of the nominal concentration applied will be bound to the Perspex recording chamber we use (Börjesson et al., 2008). As a result, the effective concentration of freely available fatty acids is 70% of the applied concentration. To allow for comparison with previous studies, we report the effective concentrations throughout the paper.

### Molecular biology

Human KCNQ4 (GenBank accession no. NM_004700) and human KCNQ5 (GenBank accession no. NM_001160133) were used in this study. cRNA for injection was prepared from DNA using a T7 mMessage mMachine transcription kit (Invitrogen, Stockholm, Sweden), and cRNA concentrations were determined by means of spectrophotometry (NanoDrop 2000c, Thermo Scientific, Stockholm, Sweden). Mutations were introduced through site-directed mutagenesis (QuikChange II XL, with 10 XL Gold cells; Agilent Technologies, Kista, Sweden) and confirmed by sequencing at the Linköping University Core Facility.

### Two-electrode voltage clamp experiments on *Xenopus* oocytes

Individual oocytes from *Xenopus laevis* frogs were acquired either through surgical removal followed by enzymatic digestion at Linköping University, or purchased from Ecocyte Bioscience (Dortmund, Germany). The use of animals, including the performed surgery, was reviewed and approved by the regional board of ethics in Linköping, Sweden (Case no. 1941). Oocytes at developmental stages V-VI were selected for experiments and injected with 50 nL of cRNA. Each oocyte received either 2.5 ng of hK_V_7.4 RNA or 5 ng of hK_V_7.5 RNA. Co-injected oocytes received a 1:1 mix of hK_V_7.4 cRNA (2.5 ng) and hK_V_7.5 cRNA (2.5 ng). Injected oocytes were incubated at 8°C or 16°C for 2-4 days prior to electrophysiological experiments.

Two-electrode voltage clamp recordings were performed with a Dagan CA-1B amplifier system (Dagan, MN, USA). Whole-cell K^+^ currents were sampled using Clampex (Molecular devices, San Jose, CA, USA) at 5 kHz and filtered at 500 Hz. For most experiments, the holding potential was set to -80 mV. If experimental conditions allowed for channel opening at -80 mV, the holding potential was set to -100 mV. Current/voltage relationships were recorded using voltage-step protocols prior to and after the application of test compounds. Activation pulses were generated in incremental depolarizing steps of 10 mV, from -100 mV to +60 mV for hK_V_7.4, from -90 mV to 0 mV for hK_V_7.5, and from -120 mV to +60 mV for hK_V_7.4/hK_V_7.5 co-injection experiments. The duration of the activation pulse was 2 seconds for hK_V_7.4 and hK_V_7.4/7.5, and 3 seconds for hK_V_7.5. The tail voltage was set to -30 mV for all protocols and lasted for 1 second. All experiments were carried out at room temperature (approx. 20°C). The extracellular recording solution consisted of (in mM): 88 NaCl, 1 KCl, 0.4 CaCl_2_, 0.8 MgCl_2_ and 15 HEPES. pH was adjusted to 7.4 by addition of NaOH. When experiments were conducted at a lower or higher pH, the pH was adjusted the same day as experiments by the addition of HCl or NaOH. Recording solution containing test compounds was applied extracellularly to the recording chamber during an application protocol (comprised of repeated depolarizing steps every 10 seconds to 0 mV for hK_V_7.4 and hK_V_7.4/hK_V_7.5 co-injected cells, or to -30 mV for hK_V_7.5) until steady-state effects were observed. Solutions containing PUFAs or PUFA analogues were applied directly and manually to the recording chamber via a syringe. A minimum volume of 2 mL was applied to guarantee the replacement of the preceding solution in the recording chamber. The recording chamber was thoroughly cleaned between cells with 99.5% ethanol.

### Electrophysiological analysis

GraphPad Prism 8 software (GraphPad Software Inc., Ca, USA) was used for data analysis. The voltage-dependence of hK_V_7 channels was approximated by plotting the immediate tail currents (recorded upon stepping to the tail voltage) against the preceding test voltages. Data were fitted with a Boltzmann function, generating a G(V) (conductance *versus* voltage) curve:

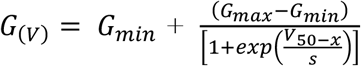

where G_min_ is the minimum conductance, G_max_ is the maximum conductance, V_50_ is the midpoint of the curve (i.e, the voltage determined by the fit required to reach half of G_max_) and s is the slope of the curve. The difference between V_50_ under control settings and under test settings for each oocyte (the ΔV_50_) was used to quantify shifts in the voltage-dependence of channel opening evoked by test compounds. The relative difference between G_max_ under control settings and under test settings for each oocyte (the ΔG_max_) was used to quantify changes in the maximum conductance evoked by test compounds.

To determine the concentration dependence or the pH dependence of ΔV_50_, the following concentration-response function was used:

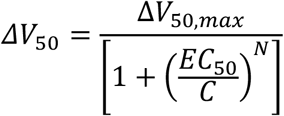

where ΔV_50,max_ is the maximum shift in V_50_, C is the concentration of the test compound, EC_50_ is the concentration of a given test compound or the concentration of H^+^ required to reach 50% of the maximum effect, and N is the Hill coefficient (set to 1 or -1). For studying pH dependence, the values of C were determined with asymptotic 95% confidence intervals in GraphPad Prism 8 and subsequently log-transformed to acquire the apparent pKa values.

### *In silico* analysis

The amino acid sequences for hK_V_7 channels were acquired from UniProt (accession codes for hK_V_7.1-hK_V_7.5 are P51787, O43525, O43526, P56696 and Q9NR82, respectively) and aligned using Clustal Omega (Sievers and Higgins, 2018). The cryo-EM resolved structure of the hK_V_7.4 channel (PDB accession code: 7BYL; (Li et al., 2021)) was visualized using The Protein Imager (Tomasello et al., 2020).

### Statistical analysis

Average values are expressed as mean ± SEM. When comparing two groups, a Student’s *t* test was performed. One sample *t* test was used to compare an effect to a hypothetical effect of 0 (for ΔV_50_) and 1 for (ΔG_max_). When comparing multiple groups, a one-way ANOVA was performed, followed by Dunnett’s multiple comparison test when comparing to a single reference group. A *P* value < 0.05 was considered statistically significant. All statistical analyses were carried out in GraphPad Prism 8.

## Acknowledgements

We thank Dr H. Peter Larsson, University of Miami, and Dr Fredrik Elinder, Linköping University, for comments on the manuscript. We thank Louise Abrahamsson for her contribution to experiments during her time as visiting scholar. The clones for human K_V_7.4 and K_V_7.5 were kind gifts from Dr. Nicole Schmitt at University of Copenhagen. This project has received funding from the Swedish Society for Medical Research and the Swedish Research Council (2017-02040).

---

**Figure 1 – Figure Supplement 1.**
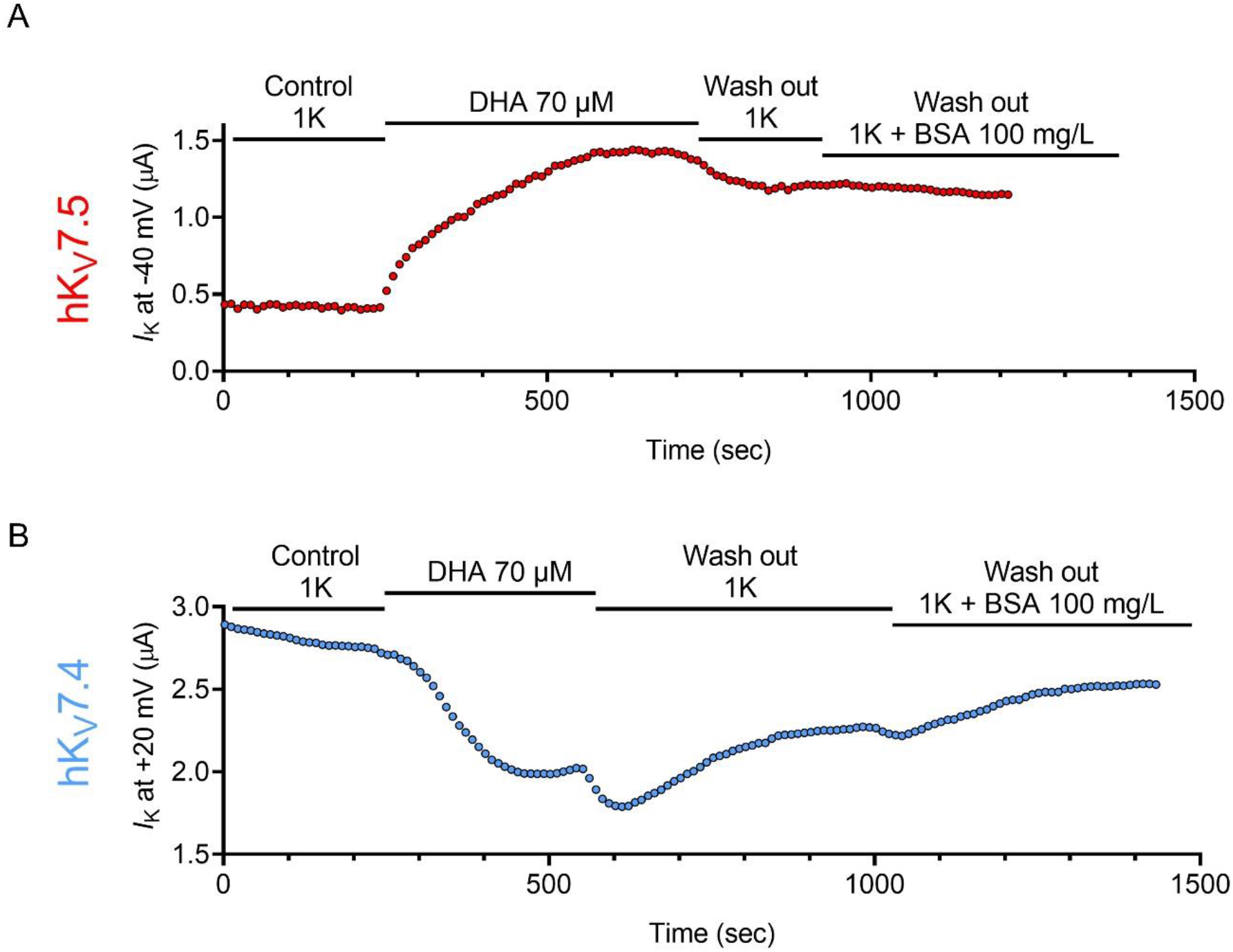
The DHA response has a rapid onset for both hK_V_7.5 and hK_V_7.4. Representative data showing wash-in and wash-out of 70 µM DHA on hK_V_7.5 (A) and hK_V_7.4 (B). Data points represent the current amplitude recorded following a test voltage to -40 mV (for hK_V_7.5) or +20 mV (for hK_V_7.4). The onset of DHA induced activation of the hK_V_7.5 channel was quick, reaching a stable level of enhanced current within 5 minutes. Neither standard recording solution nor recording solution supplemented with BSA completely removed the DHA effect, and the enhanced current amplitude remained at almost 3-fold that of the baseline current amplitude. The onset of DHA induced inhibition of the hK_V_7.4 channel was also quick, reaching a stable level of reduced current amplitude within 4.5 min. While not fully reversible, the DHA effect was reduced following re-perfusion with standard recording solution. Recording solution supplemented with 100 mg/mL BSA also reduced the inhibitory effect, with the current amplitude reaching approximately 89% of baseline current amplitude. Related to Figure 1.

